# An archaeal histone-like protein regulates gene expression in response to salt stress

**DOI:** 10.1101/2021.10.14.464415

**Authors:** Saaz Sakrikar, Amy Schmid

## Abstract

Histones, ubiquitous in eukaryotes as DNA-packing proteins, find their evolutionary origins in archaea. Unlike the characterized histone proteins of a number of methanogenic and themophilic archaea, previous research indicated that HpyA, the sole histone encoded in the model halophile *Halobacterium salinarum*, is not involved in DNA packaging. Instead, it was found to have widespread but subtle effects on gene expression and to maintain wild type cell morphology; however, its precise function remains unclear. Here we use quantitative phenotyping, genetics, and functional genomic to investigate HpyA function. These experiments revealed that HpyA is important for growth and rod-shaped morphology in reduced salinity. HpyA preferentially binds DNA at discrete genomic sites under low salt to regulate expression of ion uptake, particularly iron. HpyA also globally but indirectly activates other ion uptake and nucleotide biosynthesis pathways in a salt-dependent manner. Taken together, these results demonstrate an alternative function for an archaeal histone-like protein as a transcriptional regulator, with its function tuned to the physiological stressors of the hypersaline environment.

## INTRODUCTION

Phylogenetic analysis has shown that the histone fold domain originated in the Archaea (1-3). Histone proteins play a vital role in genome compaction and regulation of gene expression in eukaryotes (4). The four core eukaryotic histones (H3, H4, H2A, H2B) share a histone fold domain, which is involved in histone dimerization and DNA-binding (5-7). Proteins containing the histone fold are present in all known major archaeal lineages (7). Archaeal histone-like proteins have been most extensively characterized in species representing the euryarchaeal superphylum, with most work so far focusing on *Methanothermus fervidus* (8) and *Thermococcus kodakarensis* (9-11). From *in vitro* and *in vivo* studies, including structural analyses (12,13) and nuclease digestion (14), it was interpreted that archaeal histones function similarly to those of eukaryotes. These histones appeared to act as the major chromatin protein by forming extended polymeric structures that wrap DNA in multiples of 30-60 bp. Like eukaryotic histones, in some species archaeal histones have been shown to influence global transcription levels by hindering initiation (15) or elongation (16). Archaeal histones can also inhibit the binding of site-specific transcription factors (TFs) through competition (17). These studies led to the current prevailing hypothesis that archaeal histone function largely resembles that of eukaryotes in terms of genome compaction and gene expression, although some have noted key differences (7,18).

Recent evidence in other model systems challenge this hypothesis. For example, a deletion mutant of the sole histone of *Methanosarcina mazei* was viable, but exhibited reduced growth when exposed to radiation (19). Phylogenetics and molecular dynamics simulations in other model methanogens that encode multiple histone variants suggest various functions in the chromatin environment (20). Previous work from our group demonstrated an alternative regulatory function for HpyA, the sole histone of the hypersaline-adapted species *Halobacterium salinarum* (21). Like in *M. mazei*, HpyA is dispensable for cell viability. Unlike in *M. mazei* histone, HpyA is important for maintaining wild type gene expression and cell shape under optimum growth conditions. HpyA protein levels were too low to facilitate genome-wide DNA compaction (21). Phylogenetic and proteomics evidence in hyperthermophiles suggests that chromatin compaction allows DNA stability to prevent unwanted transcription by promoter melting at high temperature (22). Histone point mutants that cannot compact DNA exhibit differential expression of specific genomic regions (11). Together these findings challenge a predominant hypothesis, instead suggesting diverse histone functions across archaeal lineages selected for by the diverse and sometimes extreme environments of archaea. However, the function of histone-like proteins in hypersaline-adapted archaea remains understudied relative to other archaeal lineages.

Halophilic archaea have adapted to survive extreme osmotic pressure (up to 5M NaCl) in their natural salt lake environments by counterbalancing with up to 4M potassium ions in the cytoplasm (23). Due to the resultant highly ionic cytoplasm, haloarchaeal proteins have evolved a negatively charged surface (24), including the histone protein. This is in contrast to all other known species, where the positively charged surface of histones facilitates DNA-histone interactions (3,21). It has previously been observed *in vitro* that naked DNA under high salinity tends to spontaneously form structures similar to the beads-on-a-string observed with histone-bound DNA (25), calling into question the need for protein-based genome compaction. Based on these results and given the unusual chemistry of the haloarchaeal saturated salt cytoplasm, here we hypothesize that the non-canonical function of HpyA in gene regulation is linked to the unique hypersaline cytoplasmic environment of *Hbt. salinarum*.

We tested this hypothesis using a battery of *in vivo* quantitative phenotyping and functional genomics assays. Growth rate and cell morphology in low sodium was impaired in the Δ*hpyA* deletion strain, confirming a link between the gene and salt concentration. Protein-DNA binding assays (ChIP-seq) revealed reproducible, salt-dependent, genome-wide binding of HpyA at nearly 60 discrete sites -- a binding pattern too infrequent to coat or compact the genome. However, the high prevalence of binding within gene bodies suggests that the mechanism of regulation differs substantially from that of canonical TFs. Integration of DNA binding with transcriptomics data revealed direct regulation of iron uptake by HpyA. Global, indirect regulation of transport of other ions, biosynthesis of purines, and DNA replication and repair was also observed. Together, these results suggest that HpyA functions as a specific, direct transcriptional regulator of metal ion balance. HpyA thereby maintains growth rate and rod-shaped cell morphology during hypo-osmotic stress.

## MATERIALS AND METHODS

### Strains, media and general culturing

Strains used in this study have been described in Dulmage et al 2015 (21), summarized in **Table S1**. All strains were constructed from a *Halobacterium salinarum* NRC-1 background with *ura3* (encodes uracil biosynthesis functions) gene deleted to enable uracil counterselection (26). Growth assays were carried out using strain MDK407 (Δ*ura3*) as the parent strain (control, referred to here as wild type, or WT) and KAD100 (Δ*ura3*Δ*hpyA*) as the Δ*hpyA* deletion strain.

For ChIP-seq, strains carrying the *hpyA* gene tagged at its C-terminus with the HA epitope were used (21). The control strain was AKS134 (*ΔhpyA* deletion carrying the empty vector pMTFCHA). The experimental strain was KAD128, which contained the pKAD17 plasmid expressing HpyA-HA driven by its native promoter (primers and plasmids given in **Table S1**). pKAD17 was generated by: (a) insertion of *hpyA* into the pMTF-cHA plasmid upstream of the HA tag sequence (between the NdeI and HindIII restriction sites); and (b) replacement by isothermal ligation of the *P*_fdx_ promoter of the plasmid with the *P*_rpa200_ native promoter sequence of HpyA at the KpnI site.

The media used for all experiments was *Hbt. salinarum* complete media (CM) containing 250g/L NaCl, 20 g/L MgSO4•7H_2_O, 3g/L trisodium citrate dihydrate, 2g/L KCl, 10g/L Bacteriological peptone (Oxoid). pH was adjusted to 6.8. Media were supplemented with 50 µg/mL uracil to compensate for the uracil auxotrophy of Δ*ura3* parent and derivative strains. Reduced salt media was made identically except for NaCl, which was reduced to 199 g/L (3.4 M). For plasmid strains, 1 µg/mL mevinolin (AG Scientific) was added to liquid medium and 2.5 µg/mL to solid media to maintain selective pressure on the plasmid.

Cells were routinely streaked fresh from frozen stock onto solid medium. Individual colonies were picked from plates and inoculated into 5 mL CM (with additives when necessary) and allowed to grow for approximately 4 days at 42°C in a shaking incubator until stationary phase was reached. These starter cultures were diluted by sub-culturing to OD600 ∼ 0.02 into 50 mL of media indicated in the figures and grown until harvesting as described below.

### Growth and microscopy

For growth curve phenotyping, 9 biological replicates of Δ*ura3* (MDK407) and Δ*hpyA* (KAD100) strains were cultured in 125 mL flasks at 42° C in a shaking incubator. Optical density (OD) measurements were taken at time zero, then at 3-4 hour intervals following the initial lag phase of ∼12 hours. Raw growth data are provided in **Table S2**. Resultant growth curves were fit by logistic regression to calculate instantaneous growth rate (µ_max_) using the R package grofit (27). The code for analysis and visualization of these growth data are contained in https://github.com/amyschmid/HpyA_codes.

For microscopy, cultures of Δ*ura3*, Δ*hpyA*, and Δ*hpyA* / pKAD17 (strain KAD128) were each grown to mid-exponential phase. 8 µl aliquots were placed on a thin, flat, agarose pad impregnated with 4.3M NaCl as described (28). Cells were imaged at 100X using a Zeiss Axio Scope A1 microscope with a Pixelink PL-E421M camera. Images were analyzed for circularity using the MicrobeJ package within the ImageJ software (29). Given that circularity distributions were skewed, adjusted bootstrap percentile corrected 95% confidence intervals were calculated by 1,000-fold ordinary non-parametric bootstrap resampling of the median with replacement. The boot() package in the R coding environment was used for these calculations.

### ChIP-seq experiments

One biological replicate colony of AKS134 (Empty vector control) and four replicates of KAD128 (expressing HpyA-HA) were cultured as described above. The 50mL cultures were grown in 125mL flasks and their growth was monitored by OD600 until the time for harvesting (exponential phase: 36-50 hours, OD∼0.2-0.35, growth rate ∼ 0.032 hr^-1^; stationary phase: ∼70-140 hours, OD∼1.4-1.7, growth rate ∼0.017 hr^-1^). Strains were PCR-checked for the presence of the plasmid expressing *hpyA*-HA prior to each experiment (see **Table S1** for primers).

Harvested cells (45mL) were immediately cross-linked using 1.4mL 37% formaldehyde (final concentration = 1% v/v) and immuoprecipitated as described in Wilbanks et al, 2012 (30), with certain modifications to the protocol: the cross-linking reaction was allowed to proceed for 20 minutes, and cell pellets were resupended in 800 μL lysis buffer. Resulting DNA was extracted with Phenol:Chloroform:Isoamyl alcohol (25:24:1) and then ethanol precipitation. Library preparation and single-end sequencing was carried out by the Duke Center for Genomic and Computational Biology Sequencing and Genomic Technologies core facility using the Illumina HiSeq4000 instrument.

### Analysis of ChIP-seq data

Gzipped FastQ files (Accession: PRJNA703048, GEO: GSE182514) were analyzed using FastQC software. Information provided as input included read sequence quality, length distribution, and presence of adapters. Adapters were trimmed from the reads using Trim Galore!, and these trimmed sequences were aligned to the *Hbt. salinarum* NRC-1 genome (RefSeq ID GCF_000006805.1, assembly ID ASM680v1) to generate a SAM file using Bowtie2 with default parameters. End-to-end alignment was suitable for trimmed reads (31). FastQC and Trim Galore! are available online at http://www.bioinformatics.babraham.ac.uk/projects/ (2015 version). The SAM files were converted to binary (BAM), sorted and indexed using SAMtools (32). Sorted BAM files were used for peak calling. WIG files for easy visualization were also generated using SAMtools, with coverage recorded every 10 bp. All code used to analyze ChIP-seq data are available in File S1 at https://github.com/amyschmid/HpyA_codes.

The sorted BAM files were used for peak-calling with MACS2 (33) version 2.1.1 callpeak function. Parameters were: nomodel, qval=0.05 cutoff. Called peaks were combined across replicates using the *multiBedIntersect* function of the bedtools package (34). Only peaks detected in at least two biological replicate experiments were kept in downstream analyses. Genes within 500 bp of these reproducible peaks were annotated using the IRanges package in R (35). Resultant peaks were then manually curated to remove the following: (a) false positives caused by local variability in input control sequencing read depth; (b) local duplications and deletions associated with transposases and integrases; (c) one peak that was also detected in the HA tag-alone input control; (d) peaks located nearby redundant genes. Details of the code and dependencies for the entire workflow for peak calling and visualization are noted in the github repository https://github.com/amyschmid/HpyA_codes.

Resultant peaks were then classified based on their genomic locations and context to tabulate the results given in Figure 4 and the corresponding text (details in **Table S3**). Promoters were defined as the region from 500bp upstream of the translation start site [many halophile transcripts are leaderless (36)]. To classify binding locations “genic” or “promoter”, the number of bp in the overlap between the ChIP-seq peak chromosomal coordinates and the genomic feature was calculated. If the peak overlapped both a genic and a promoter feature, the peak was classified as located within the feature with the largest overalp. Both features were counted in the case of ties. The code used to make this classification is in https://github.com/amyschmid/HpyA_codes. Operons were computationally predicted using the Operon-Mapper tool (37) and integrated with empirical predictions from Koide et al (36). Classification of TrmB binding locations are given directly in reference (38) and significance of enrichment was computed using the hypergeometric test in R. Classification and computation of enrichment *p*-values for RosR binding locations [from reference (39)] were computed using BEDtools “fisher” function (40).

**Figure 1:**
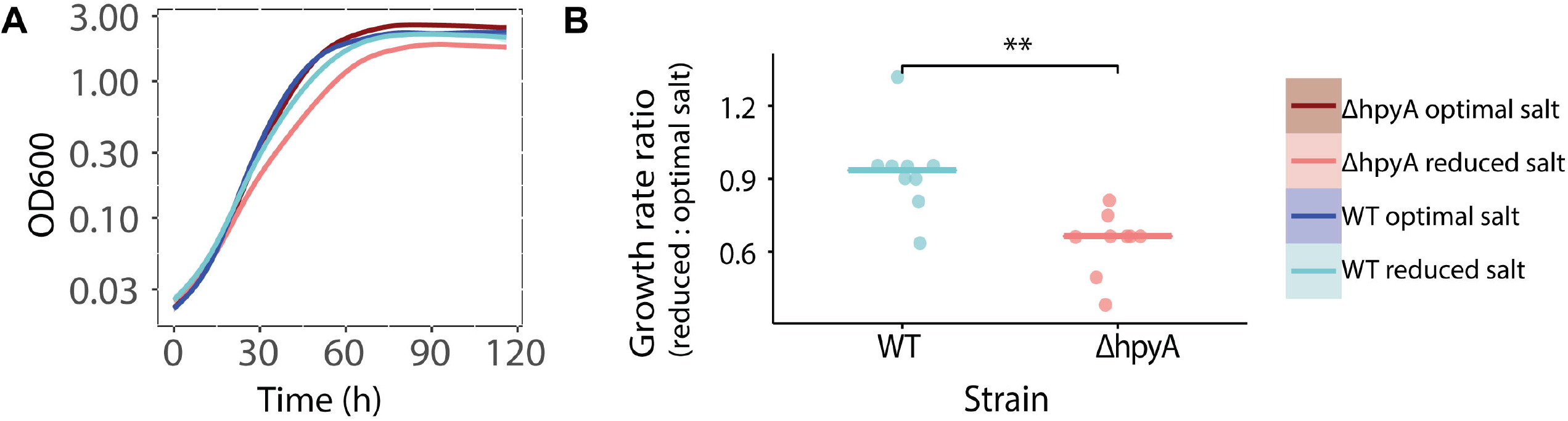
The Δ*hpyA* strain is impaired for growth under reduced salt conditions. (**A**) Spline-smoothed growth curves for the Δ*ura3* parent (“WT”, blue curves) compared to the Δ*hpyA* mutant (red curves) under optimal salt (dark colors) and reduced salt (light colors). For each strain under each condition, the mean of 9 biological replicate growth curves is shown with surrounding shaded region representing the 99% confidence interval (CI, in some curves, shading is not visible because the mean line and CI overlap). (**B**) Dot plots of relative maximum instantaneous growth rate (µ_max_) for each of the WT and Δ*hpyA* strains. Each dot represents one of nine biological replicate trials measuring the µ_max_ for each strain under reduced salt compared with its own growth in optimal conditions. Horizontal bars represent the median of each distribution.

**Figure 2:**
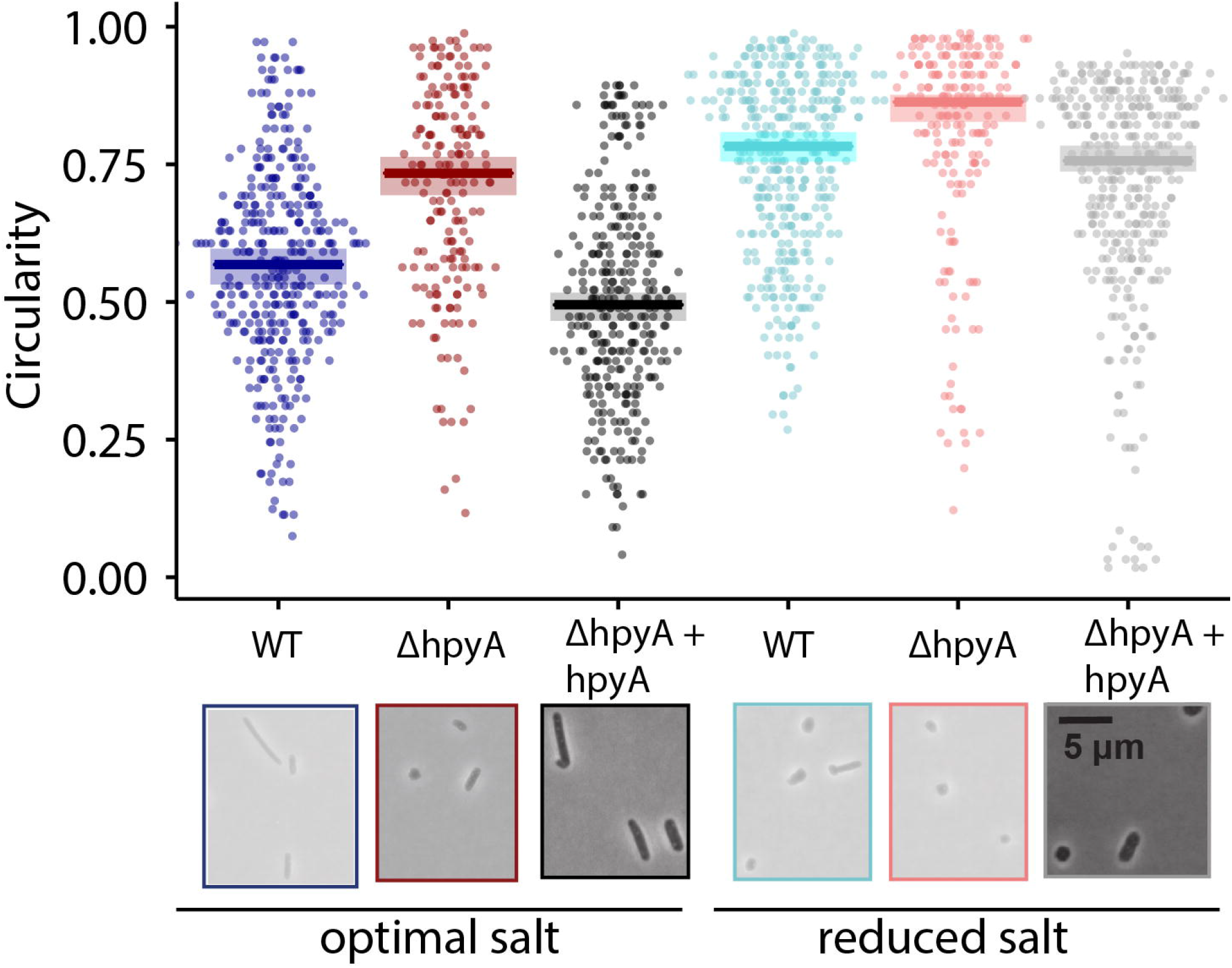
Circularity of *Hbt. salinarum* increases when *hpyA* is deleted under reduced salt. In dot plot, dots represent circularity measurements of individual cells. Horizontal bars are the median of the distribution in each strain under each condition. Shaded regions represent the 95% bias-corrected confidence interval from bootstrap resampling (see Methods). Below, representative micrograph images are shown for cells of WT, Δ*hpyA*, and complemented strain (*ΔhpyA + hpyA*, i.e. pKAD103, **Table S2**) cells in optimal and reduced salt media. Scale bar is 5 µm and consistent across images. Colors are as in Figure 1. Number of cells counted: WT in optimal salt, n = 363; WT in reduced salt, n = 383; Δ*hpyA* in optimal salt, n = 188; Δ*hpyA* in reduced salt, 187; complemented strain in optimal salt, n = 313; complemented strain in reduced salt, n = 360.

**Figure 3:**
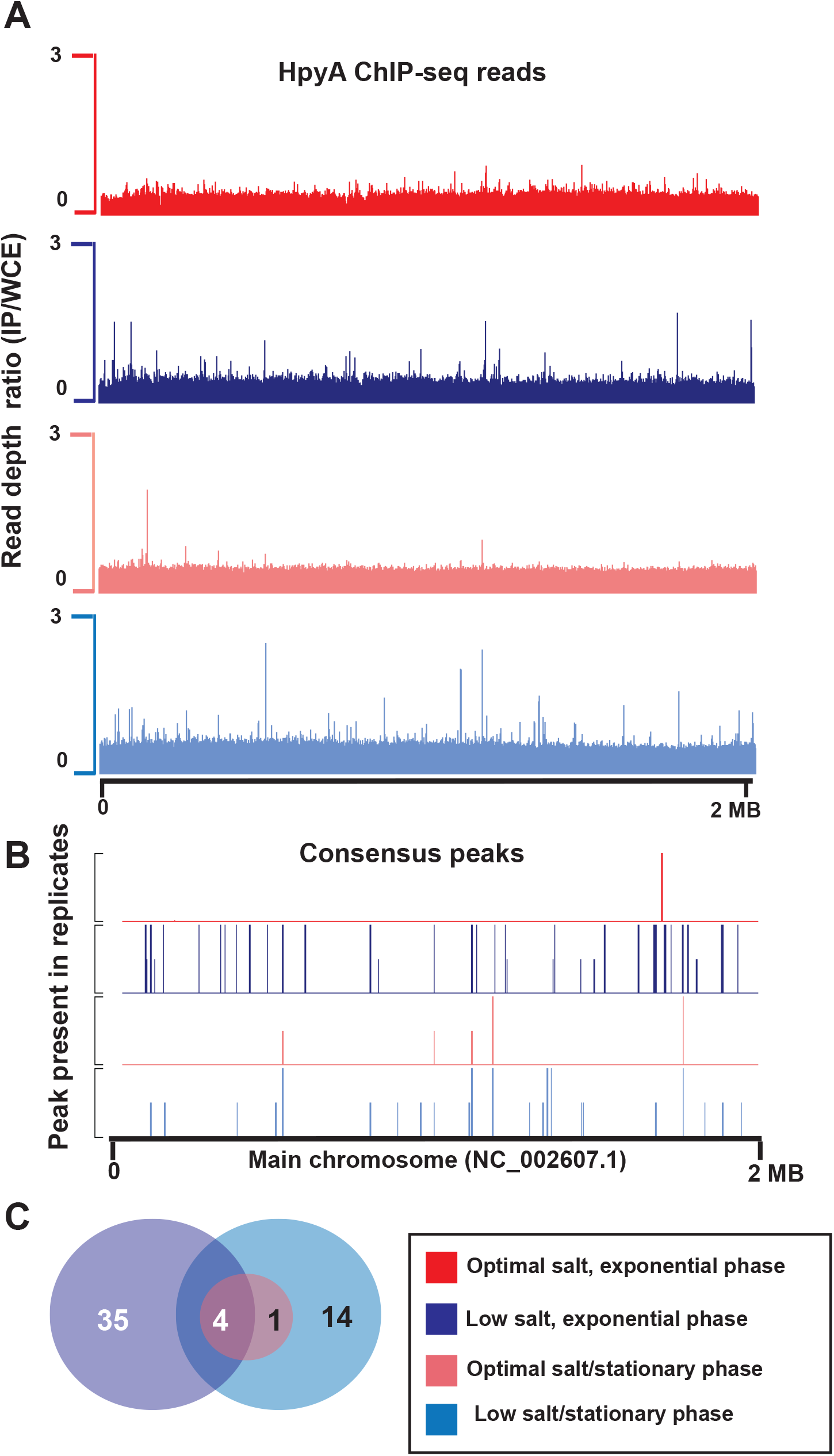
ChIP-seq of HpyA shows salt and growth phase dependent binding patterns. (**A**) Chromosome-wide binding pattern (measured as read-depth of IP/Input) of HpyA-HA in optimal salt and exponential growth phase (red), optimal salt and stationary phase (pink), reduced salt and exponential phase (dark blue), reduced salt and stationary phase (light blue). (**B**) Reproducible peaks detected across at least 2 of 4 biological replicates for each condition – shorter peaks represent those found in 2 replicates only, while peak at full heights were detected in at least 3 replicates for that particular condition. Note that peaks shown in tag-alone control have been removed from the other conditions and from further analysis. (**C**) Venn diagram indicating the number of peaks detected in the different conditions. Circles are not scaled by number of peaks.

**Figure 4:**
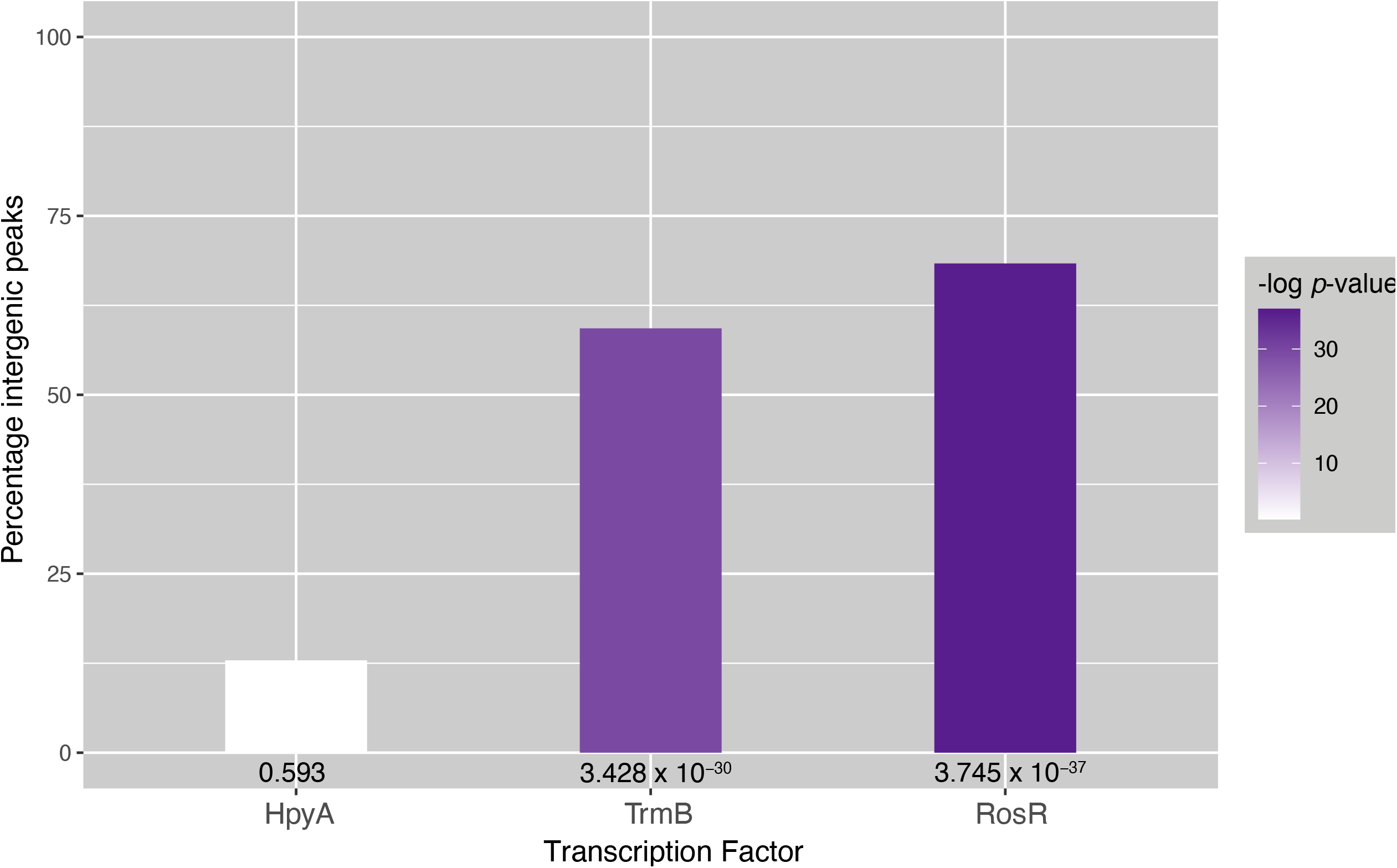
HpyA binds without preference for coding vs non-coding regions. Height of the bar graph corresponds to percentage of ChIP-seq peaks in non-coding (intergenic, promoter) regions of the genome. Color of the bars are shaded by negative log10 *p*-value of enrichment of peak locations in promoter regions (see scale at right). Actual *p*-values of enrichment calculated by hypergeometric test for each TF are also written below each bar. HpyA binding locations (left) are compared with those for characterized TFs TrmB and RosR in *Hbt. salinarum* (Hbt) (38,39).

### RNA-seq experiments

Six biological replicate cultures of strains MDK407 (parent) and KAD100 (Δ*hpyA*) were cultivated as described above in either optimal salt (4.2M NaCl) or low salt (3.4M NaCl) media. Growth was monitored using OD600 until harvesting (exponential phase was defined as: ∼31-34 hours of growth, OD∼0.1-0.4 depending on the strain and medium).

A 4.2mL aliquot of each culture was removed and centrifuged for 30s at 21,000 x g in an Eppendorf tabletop centrifuge. The supernatant was discarded and the cell pellet was immediately plunged into liquid nitrogen and stored 1-7 days at −80° C. Extraction of RNA from these pellets was carried out using the Agilent Absolutely RNA Miniprep kit following the manufacturer’s protocol, with an extended on-column DNase incubation of 45-60 min. Resultant RNA samples were checked for: (a) genomic DNA contamination using PCR with 200ng input RNA and 35 amplification cycles using primers listed in **Table S1**; (b) concentration using 260/280 nm ratio in a Nanodrop spectrophotometer; (c) quality using the Agilent Bioanalyzer RNA Nano 6000 chip (RNA Integrity Number (RIN) > 9.0). For each strain and condition, rRNA was removed from 3 replicates with NEBNext Bacteria rRNA Depletion Kit (New England Biolabs), while the other 3 were treated with NEBNext Depletion Core Reagent Set using custom probes targeted to *Haloferax volcanii* rRNA (Martinez-Pastor and Sakrikar, unpublished). These custom probes were designed using the NEBNext Custom RNA Depletion Design Tool (https://depletion-design.neb.com/). rRNA depletion was verified using the Bioanalyzer RNA chip. The NEBNext Ultra II Directional RNA Library Prep Kit for Illumina was used for preparing sequencing libraries, and cDNA libraries were quality-checked using the High-Sensitivity DNA Bioanalyzer chip. Paired-end sequencing was carried out at the Duke Center for Genomic and Computational Biology Sequencing core facility using the Novaseq6000 instrument (Illumina).

### RNA-seq data analysis

For analysis of sequencing data, paired FastQ files were trimmed and checked for quality using Trim Galore! (http://www.bioinformatics.babraham.ac.uk/projects/) and aligned to the genome using Bowtie2 (31). SAMtools was used to generate, sort, and index BAM files (32). The count function of HTSeq (41) was used to create a file assigning the number of reads to each gene (see **File S1** within the Github repository for details). Outlier samples were removed from further analysis using Strong PCA (42) (https://github.com/amyschmid/HpyA_codes). The R package DESeq2 (43) was used to normalize counts and batch correct across replicates for each strain and genotype (using DEseq2 default parameters). Significant differential gene expression analysis using DESeq2 applied three pairwise contrasts: Δ*hpyA* vs WT in optimal salt, Δ*hpyA* vs WT in reduced salt, and reduced vs optimal salt in a WT background. For each contrast, reproducibility and quality was checked across replicates using dispersion, MA, and volcano plots. For each contrast, Benjamini-Hochberg (BH) adjusted (44) Wald test *p* < 0.05 (default within DESeq2) was used as the criterion for significant differential expression (results in **Table S4**).

Averaged normalized counts across biological replicates for each strain and stress treatment were then mean and variance standardized and subjected to Kmeans clustering using the factoextra package in R, which also determines the best value for K (45). Resultant gene clusters were then subclustered using Kmeans and visualized using ggplot2 (46) and pheatmap (47) functions in R (https://github.com/amyschmid/HpyA_codes). This clustering procedure was carried out twice, once with genes differentially expressed in both reduced and optimal salt, and then excluding genes differentially expressed in optimal salt. Results of the clustering are given in **Table S5**. For analysis of gene functional enrichments, the hypergeometric test *p*-value of enrichment for differentially expressed genes was calculated. Resultant *p*-values were BH-corrected for multiple hypothesis testing. The archaeal Clusters of Orthologous Genes (arCOG) functional ontology was used for functional assignments (48), results are listed in **Table S6**.

## RESULTS

### HpyA is important for wild type growth and morphology in low salinity stress conditions

To test the hypothesis that HpyA plays a role in salt stress, we compared the growth rate of *Hbt. salinarum* Δ*ura3* (parent strain, hereafter referred to as wild type or “WT”) to Δ*hpyA* cells in rich complete medium with salt concentrations supporting optimal growth (CM, 4.2M NaCl) and CM with reduced salt (3.4M NaCl). As expected from previous observations (21), instantaneous growth rate (µ_max_) under optimal salt of the WT strain was statistically indistinguishable from that of Δ*hpyA* **(Fig. 1A, Table S2, Fig. S1**). Reduced salt slows the instantaneous growth rate (µ_max_) of WT cultures to 89% of that in standard conditions. In contrast, *ΔhpyA* cultures show significant growth impairment in reduced salt relative to WT, growing at 67% of their standard rate. (**Fig. 1B;** unpaired two-sample *t*-test *p* < 0.008).

Cell morphology of *Hbt. salinarum* changes from rod-shaped to circular in the presence of low salt (49,50). Our previous work demonstrated that the Δ*hpyA* strain exhibits similar circularity in standard conditions (21). To further test the hypothesis that HpyA plays a role in the salt stress response, we used phase contrast microscopy to visualize the combined effects of reduced salt and *hpyA* deletion on cell shape. From the images, we quantified circularity of individual cells (where 1 indicates a perfectly circular cell). In media containing optimal salt concentrations, WT cells are primarily rod-shaped, whereas the Δ*hpyA* cells are significantly rounder (**Fig. 2**, non-parameteric bootstrapped 95% confidence intervals of the medians of these distributions do not overlap, see Methods). In reduced salt, WT cell morphology was more circular: the distribution was indistinguishable from that of Δ*hpyA* in optimal salt. Δ*hpyA* morphology in reduced salt was the most circular of all strain-by-genotype combinations, indicating that this strain’s morphology is strongly impacted by reduced salt.

Growth and morphology defects are significantly complemented by expression of *hpyA* from its native promoter *in trans* on a plasmid (Δ*hpyA* + *hpyA*-HA, **Fig. 2**). Whole genome resequencing of the Δ*hpyA* strain also demonstrated that: (a) second site suppressor mutations were absent; and (b) deletion of *hpyA* was complete through all chromosomal copies (**Table S1**). *Hbt. salinarum* is highly polyploid (51), which necessitates validation that all gene copies have been deleted. These results indicate that Δ*hpyA* phenotypes are solely attributable to the deletion of *hpyA*.

Because cell shape differences can lead to alterations in light scattering in a spectrophotometer (52), as a control, we calculated CFU/mL by dilution plate counting. We found that OD600 measurements were well correlated with CFU counts for both strains and media preparations. In optimal salt, WT cultures CFU to OD Spearman correlation was ρ = 0.81 (*p* = 0.00081), Δ*hpyA* ρ = 0.64 (*p* = 0.015). In reduced salt, WT correlation was ρ = 0.93 (*p* < 2.2 × 10^−16^), Δ*hpyA* ρ = 0.78 (*p* = 0.001; **Fig. S2**). This indicates that the Δ*hpyA* growth defect observed in reduced salt (as measured by optical density) is due to differences in growth and not an artefact of the shape change. Taken together, these batch culture (**Fig. 1)** and single cell microscopy (**Fig. 2)** quantitative phenotype data suggest that HpyA is important for maintaining wild type morphology and growth in response to hypo-osmotic salt stress.

### HpyA binds genome-wide in a salt-specific manner

To determine which genes are potential targets of transcriptional regulation by HpyA, we performed genome-wide DNA binding location analysis using chromatin immunoprecipitation coupled to sequencing (ChIP-seq). For this purpose, we generated an Δ*hpyA* strain expressing *in trans* HpyA translationally fused at its C-terminus to the hemagglutinin (HA) epitope tag. This fusion construct was driven by its native promoter (see Methods and **Table S1** for strain details). As described above, expression of HpyA-HA *in trans* complemented the circularity defect of Δ*hpyA*, demonstrating that HA tag and plasmid-based expression does not interfere with wild type function of HpyA (**Fig. 2)**. Based on the Δ*hpyA* phenotypes observed (**Figs. 1 & 2**), the ChIP-seq experiments were performed at both physiological and reduced salt concentrations in both mid-exponential and stationary phase. HpyA binding was enriched relative to the background input control at a total of 59 discrete genomic locations (ChIP-seq peaks) across all conditions tested (**Table S3**). These 59 peaks were consistently detected in reduced salt across growth phases and biological replicate experiments **(Figure 3A and B**), but only 5 of these peaks remained bound in optimal salt conditions. Of the low salt peaks, 35 were detected exclusively during exponential growth phase, 14 exclusively during stationary phase, and 9 across both growth phases (**Figure 3C**). Because HpyA binds DNA primarily under low salt conditions, these results corroborate the growth and morphological impairments of Δ*hpyA* cells observed in early log phase under reduced salt conditions (**Figure 2)**.

HpyA binding peaks were located nearby 86 genes (within the gene coding region or 500 bp upstream of the gene start in the promoter / non-coding region). Few of the HpyA binding sites are located within non-coding or promoter regions of the genome (12.9%, *p* = 0.593). In contrast, other previously characterized *Hbt. salinarum* TFs bind in a sequence-specific manner with significant preference non-coding regions [**Fig. 4**, (38,39)]. Binding of HpyA is also not statistically enriched for binding withing gene coding regions (*p* = 0.152). This high number was as expected because, like many archaeal genomes, the *Hbt. salinarum* genome is dense with coding sequences (86%). Taken together these DNA binding results suggest that, unlike canonical histone proteins of eukaryotes and other archaeal species, HpyA binds in a salt-specific manner to a restricted set of sites genome-wide. However, unlike canonical TFs, HpyA binds apparently without preference for coding vs non-coding regions.

### HpyA functions primarily as an activator of genes encoding ion transport and metabolic proteins

Based on the quantitative phenotyping and ChIP-seq data, we reasoned that HpyA may regulate gene expression in response to salt stress. To test this hypothesis, we performed transcriptome profiling experiments in WT vs Δ*hpyA* strains in both optimal and reduced salt using RNA-seq (see **Methods**). In the WT strain, over one-third of the genes in the transcriptome were significantly differentially expressed during exponential growth phase in reduced salt compared to optimal salt conditions (*p* < 0.05; 882 genes; 37% of genome; **Table S4**). Of the 37 genes previously identified by microarray analysis (53), 22 genes were also identified as significantly differentially expressed in the current dataset. For these 22 genes, the fold-change in expression was strongly and significantly correlated across the two datasets (ρ = 0.86, *p* ρ2.2 × 10^−16^). Our results therefore recapitulate but also extend previous observations that *Hbt. salinarum* mounts a strong, reproducible, and global regulatory response to hypo-osmotic stress.

To determine the extent of HpyA’s regulatory reach, gene expression ratios (Δ*hpyA*:WT) were calculated during mid-exponential growth in optimal salt and reduced salt conditions (in two separate DEseq2 analyses, see **Methods**). A total of 168 differentially expressed genes (DEG) were detected, 143 of which were significantly altered in reduced salt and 46 in optimal salt in Δ*hpyA* vs WT (**Figure 5A, Table S4**). Of these, 121 genes were uniquely differentially expressed in response to low salt. These genes are significantly enriched for a wide variety of functions critical to maintaining cell growth and physiology in adverse conditions, especially ion transport and nucleotide metabolism (hypergeometric test *p* < 0.05 enrichment in arCOG categories(48), **Figure 5B, Table S6**). Across both optimum and reduced salt conditions, the expression of 21 genes was significantly affected by *hpyA* deletion. These genes encode predicted functions in DNA recombination, replication, and repair pathways including RadA, DNA topoisomerase VI, and RPA family proteins (**Table S4, Table S6**).

**Figure 5:**
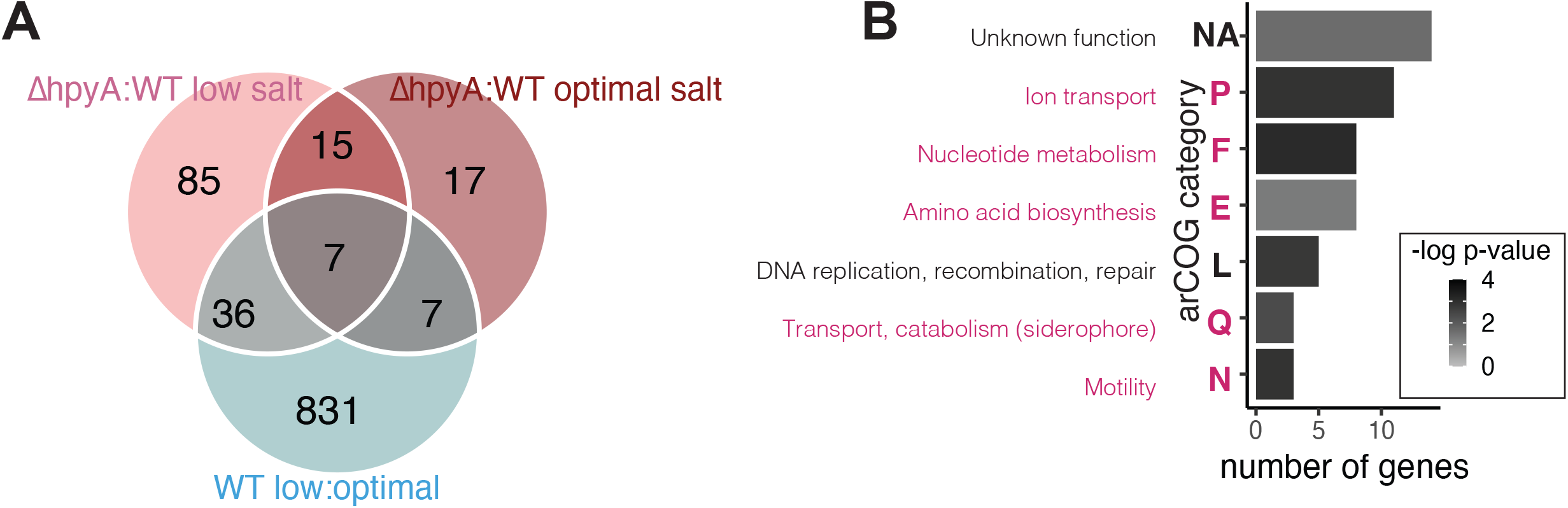
HpyA regulates gene expression in a salt-dependent manner. (**A**) Venn diagram illustrates the number of genes differentially expressed due to knockout of *hpyA* in different conditions. Genes with significant Δ*hpyA*:WT ratios in optimal salt are shown in red, genes with significant Δ*hpyA* : WT ratios low salt in pink, genes with significant low : optimal salt ratios in WT in blue. (**B**) arCOG enrichment of differentially expressed genes. X-axis shows the number of differentially expressed genes in each category that are annotated in the arCOG ontology, y-axis lists the arCOG category functions and short-hand single letter designations. Categories enriched in low salt are listed in pink text, categories enriched across conditions in black. Bars are shaded by Benjamini-Hochberg corrected (44) *p*-values of significance of enrichment according to the scale shown in the legend.

To determine the role of HpyA in the activation or repression of these genes, we performed K-means clustering analysis of normalized read count data for gene expression across the four conditions tested (Δ*hpyA* in low salt, Δ*hpyA* in optimal salt, WT in low salt, WT in optimal salt, details in Materials and Methods). We first analyzed the expression of the 21 genes that are differentially regulated the Δ*hpyA* strain in both optimal and reduced salt conditions. These 21 genes fall into 2 clear categories – 10 genes downregulated in the Δ*hpyA* strain and 11 genes upregulated (**Fig 6A, Table S5**). As noted above, genes across these two clusters are significantly enriched for DNA recombination, replication, and repair functions (8 genes). HpyA binding was detected in ChIP-seq by only one of these genes (*ssb*, encoding single-stranded DNA binding protein, **Table S7**). HpyA binding was not detected for the 20 other genes in this cluster, indicating indirect regulation by HpyA.

**Figure 6:**
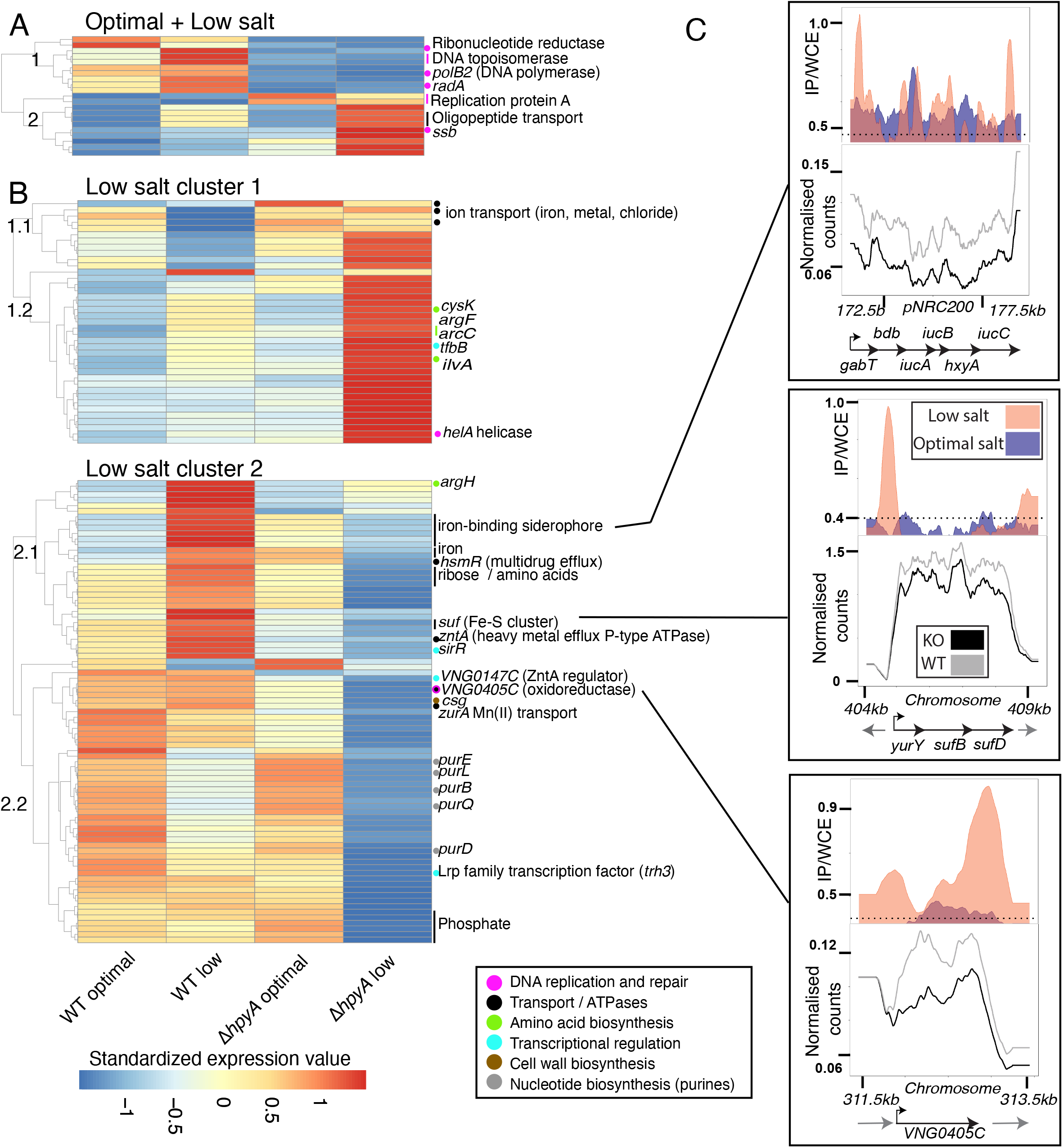
HpyA-dependent regulon shows diverse expression patterns in the conditions tested. (**A**) Clustering heatmap of genes differentially expressed in response to *hpyA* deletion across both optimal and low salt conditions. Each column corresponds to the genotype in each condition and rows represent the averaged normalized counts for each gene. Each row is self-standardized for normalization. Genes labeled with certain colors represent gene functional categories (see legend for colors). Dots next to genes represent monocistronic genes, vertical bars indicate differentially expressed operons. (**B**) Clustering heatmap of genes differentially expressed in response to *hpyA* deletion in low salt conditions alone. (**C**) Normalized reads for 3 selected direct targets of HpyA. Each box corresponds to a particular gene target indicated in the heatmap. In each panel, ChIP-seq data are shown in the top box, RNA-seq data in the middle, genomic context at bottom. ChIP-seq y-axes represent the ratio of IP to input control (whole cell extract, or WCE). RNA-seq y-axes represent read depth for WT in reduced salt (grey traces) and KO in reduced salt (black). Genonmic context images include the differentially expressed gene(s) (black arrows) and neighboring genes (grey arrows).

A separate clustering analysis of the 122 genes differentially expressed only in reduced salt in Δ*hpyA* yielded two main patterns (**Figure 6B, Table S5**). In cluster 1, genes are elevated in expression in the Δ*hpyA* background under reduced salt relative to WT, whereas cluster 2 genes are downregulated. Cluster 2 includes 64% of genes differentially expressed in low salt, suggesting HpyA functions as an activator in the majority of cases in reduced-salt conditions.

To more clearly observe the gene expression patterns and the function of differentially expressed genes, we further divided these two main clusters, resulting in a total of 4 sub-clusters (**Figure 6B, Table S5**). Subcluster 1.1 contains 11 genes whose expression pattern is downregulated in WT in reduced salt but upregulated in Δ*hpyA*. This cluster includes 3 genes predicted to encode ion transport proteins (chloride, iron, and other metals). Subcluster 1.2 contains 28 genes that are upregulated in reduced salt in WT but more heavily upregulated in reduced salt in the knockout strain. The function of genes in subcluster 1.2 are varied and not statistically enriched for a particular function. However, notable among genes in cluster 1.2 include transcription factor B (TFB), four amino acid biosynthesis genes, and HelA ATP-dependent DNA helicase (**Figure 6A, Table S5**). ChIP-seq enrichment for HpyA binding was not detected nearby any of the genes in subclusters 1.1 and 1.2, suggesting indirect regulation (**Figure 6B, Table S7**). HpyA is therefore necessary but not sufficient for repression of cluster 1 in low salt conditions.

Cluster 2 contains many genes encoding transporters (21 genes across both subclusters 2.1 and 2.2). Notably, genes encoding known metal cation transporters exhibit tight clustering with their cognate transcriptional regulators SirR and VNG0147C (54,55)(**Table S5**). Subcluster 2.1 consists of 37 genes modestly upregulated in reduced salt in the WT but strongly downregulated in reduced salt in the Δ*hpyA* mutant. Interestingly, ChIP-seq enrichment for HpyA binding was detected at 4 sites nearby genes in this subcluster (**Table S7**). Three of these 4 sites are nearby genes involved in maintenance of iron levels (**Fig. 6C**). These encode the siderophore (iron chelator) biosynthesis and transport operon, the Suf iron-sulfur cluster biosynthesis and transport system, and a putative oxidoreductase (VNG0405C). Surprisingly, the siderophore biosynthesis operon is bound at both the 5’ and 3’ ends by HpyA, which is associated with significant activation of this operon in low salt conditions (**Fig. 6C**, top panel). These results indicate that HpyA is required for direct activation of iron uptake under low sodium.

In subcluster 2.2, 46 genes are downregulated or constitutive across optimal and reduced salt in the WT, but more heavily downregulated in Δ*hpyA* in reduced salt. All 8 differentially expressed nucleotide metabolism genes are found within this subcluster, and all encode *de novo* purine biosynthesis enzymes. However, only one of the 46 genes of subcluster 2.2 is a direct target of HpyA (*VNG0161G*, encoding glutamate dehydrogenase). This suggests that HpyA regulates purine biosynthesis and other functions in this subcluster in an indirect manner.

Together, these transcriptome profiling data integrated with ChIP-seq binding locations suggest an important role for HpyA as specific, direct activator of iron uptake, and an indirect global regulator of ion transport and nucleotide biosynthesis during hypo-osmotic stress.

## DISCUSSION

Here we integrate quantitative phenotyping and functional genomics data to demonstrate that the sole histone-like protein encoded in the hypersaline adapted archaeal species *Hbt. salinarum* directly activates ion uptake transporters under hypo-osmotic stress. HpyA also functions as an indirect, global activator of genes encoding functions central to cellular physiology in low ionic strength medium. These transcriptional effects enable cells to maintain rod-shaped cellular morphology and growth in hypo-osmotic conditions.

Two of the five operons under the direct transcriptional control of HpyA encode transmembrane ABC transporters that are predicted to import iron. One operon (*VNG0524G-VNG0527C*) encodes a putative iron-sulfur (Fe-S) cluster assembly system of the Suf family. The predicted encoded proteins exhibit moderate identity to the well-characterized *E. coli* Fe-S assembly proteins SufC, SufB, and SufD (45%, 56%, and 28%, respectively (56)). The other operon (*gabT/bdb/iucABC, VNG6210-VNG6216*) encodes siderophore biosynthesis and uptake. Siderophores are high-affinity iron binding chelators that are secreted from the cell and then imported via a dedicated ABC transporter (57). In *Hbt. salinarum* and many bacteria, in addition to the ABC transporter, this operon includes a novel L-2,4-diaminobutyrate decarboxylase (DABA DC; encoded by *gabT*) and a DABA aminotransferase (encoded by *bdb*) for siderophore biosynthesis in lieu of synthesis via polyamines (57). Because amino acids are precursors for DABA biosynthesis, down-regulation of *iucABC* in the Δ*hpyA* mutant strain may also explain the indirect differential expression of amino acid biosynthesis genes during hypo-osmotic stress.

*Hbt. salinarum* is a facultative anaerobe capable of aerobic and anaerobic respiratory metabolism (58). Across the tree of life, including *Hbt. salinarum*, iron is an essential cofactor for the function of respiratory complexes in the oxygen-accepting electron transport chain (59). Because reduced salinity increases oxygen saturation in the medium, these conditions would favor aerobic respiratory metabolism over anaerobic metabolism, increasing the cellular demand for iron (60). Indeed, we observe that these iron transport systems are induced in an HpyA-dependent manner under low salt conditions (**Figure 6B**). Low levels of iron transport expression in the Δ*hpyA* strain would therefore be expected to lead to low intracellular iron levels. Low intracellular iron has also been observed previously for strains deleted for *idr2*, which encodes a DtxR family iron-dependent TF in *Hbt. salinarum*. This TF also functions as a direct activator of the *iucABC* siderophore biosynthesis and transport operon (54,61). Idr2 is a member of a complex network of TFs that regulate the response to iron imbalance (54,61) and the current study suggests that HpyA is also involved in regulation of iron uptake. This mechanism of transcriptional regulation by HpyA explains the Δ*hpyA* growth impairment observed in low sodium conditions tested here (**Fig 1**).

The remaining three operons under direct HpyA regulation encode central metabolic functions (**Table S7**). HpyA activates glutamate dehydrogenase and acyl-coA ligase enzymes, encoded by the *gdhB / alkK* operon. These enzymes control the entry of glutamate into the TCA cycle via the conversion of glutamate to 2-oxoglutarate. Glutamate is also a key precursor for biosynthesis of many metabolites, including purines and other amino acids (62,63). Direct control of this operon may explain the indirect transcriptional dysregulation of these pathways in the Δ*hpyA* mutant strain. HpyA activates an oxidoreductase gene (NAD-dependent epimerase predicted to act on nucleotide-sugar substrates; *VNG0405C*) and glycerol dehydrogenase gene and its associated operon (*VNG0161G / VNG0162G*), also encoding key components of core metabolism. The gene encoding a single-stranded DNA binding protein (*ssb*) is the only direct target predicted to be regulated by HpyA under both optimal and low salt conditions, and repressed rather than activated. Although the precise relationship between these HpyA regulatory targets and the Δ*hpyA* growth defect remains unclear, current knowledge of metabolism in *Hbt. salinarum* suggests that, in the Δ*hpyA* mutant, disruption in the levels of key metabolic intermediates (glycerol, glutamate) may contribute to the growth impairment of this strain under low salt conditions.

Dysregulation of import and/or efflux of other ions (divalent metal cations, chloride, and other transporters) in the Δ*hpyA* mutant may also explain the cell shape change in this strain (**Fig 2**). The proteinaceous surface layer (S-layer) is a key cell shape determinant of *Hbt. salinarum* (64,65). The S-layer is pliable and allows for changes in cell shape under physical pressure and low salinity (21,49). This shape change is exacerbated in Δ*hpyA* (**Fig 2**), which we hypothesize is due to dysregulation of ion transport expression. Iron has also been shown to impact cell morphology in the related haloarchaeal species *Haloferax volcanii*, although the underlying mechanism remains unknown (66). Expression of other pathways, for example, the S-layer (encoded by *csg*) and glycosylation enzymes (*VNG0140G;* **Fig 6 and Table S4**) is reduced in the Δ*hpyA* mutant under low salt conditions. However, these appear to play a more minor role in the Δ*hpyA* morphology defect given that: (a) these genes are indirect targets of HpyA regulation; and (b) overall S-layer glycosylation is unaffected in strains deleted of *hpyA* (21). Taken together, these data suggest that HpyA salt-dependent regulation of ionic balance is a major contributor to maintainance of wild type cell morphology and growth in reduced sodium environments.

Apart from these cases of direct regulation by HpyA, the majority of differentially expressed genes are located >500 bp away from HpyA binding sites. This can be explained in a number of ways. Several TFs are differentially expressed in the Δ*hpyA* strain relative to WT in low salt (**Table S4, Figure 6**). Therefore, the proximate cause of indirect differential gene expression can be inferred based on prior knowledge of the global gene regulatory network (GRN) in this organism (39,67,68). For example, the general TF, TfbB, is differentially expressed in Δ*hpyA* in low salt (cluster 1.1, **Fig 6**), and most of the genes in this cluster are indirectly regulated. TfbB is a direct regulator of several of the genes in this cluster, including *cysK* (68). *Hbt. salinarum* encodes 7 paralogs of transcription factor B (TFB) (69). Together, TFB and TATA binding protein recruit RNA polymerase to core promoters to initiate transcription (70). The TFB network in *Hbt. salinarum* is highly interconnected: for example, TfbB directly activates TfbG, which in turn regulates other genes indirectly regulated by HpyA (e.g. metal transporter *VNG1744H*, **Table S4**). Other indirect regulation by HpyA can be attributed to metal-responsive TFs. For example, SirR and VNG0147C, members of cluster 2.1, have previously been experimentally characterized as regulators of operons encoding metal transporters, specifically manganese uptake (ZurA) and the heavy metal efflux (ZntA), respectively (54,55). Aside from indirect regulation as part of a transcriptional network, we note that our data do not exclude the possibility that HpyA may function as a co-regulator, perhaps by binding DNA through interaction with another TF. Hence, we propose that HpyA may, in part, achieve its global, indirect regulatory effect via its regulation of genes encoding other TFs and/or through protein-protein interaction with other sequence-specific TFs.

In addition to transcriptional regulation of ion balance, HpyA may play other functional roles during hypo-osmotic stress. More than 40 HpyA binding sites were detected with no corresponding significant change in gene expression in the Δ*hpyA* knockout (**Table S3, S7**). HpyA prefers to bind neither coding nor non-coding genomic regions, setting it apart from characterized haloarchaeal TFs that function by canonical, sequence-specific DNA binding to promoter regions [TrmB (38) or RosR (39), **Fig. 4**]. We provide evidence of direct regulation both among targets bound in promoter and genic regions (**Table S7**). Direct regulation of expression via binding in gene bodies has also been reported for *E. coli* regulator RutR (71). Our data do not rule out that non-canonical binding modes of HpyA could also influence other aspects of the transcription cycle, including elongation or termination. Bacterial nucleoid associated proteins (NAPs) bind DNA to regulate gene expression, remodel chromatin by bending or wrapping, and/or protect the nucleoid during stress (22,72). For example, the *E. coli* transcription regulator CRP can function both as a canonical TF (site-specific gene regulation) for some genes, and as a DNA-bending chromatin remodeler at other genomic sites (22,72). These newly-discovered and expanding roles for DNA binding proteins calls for a broader perspective on the function of transcriptional regulators (22,72). Likewise, further research is needed to explore such functional roles for HpyA.

Taken together, the results presented here strongly suggest that HpyA functions as a direct activator of iron regulatory genes and a global indirect regulator of diverse pathways. This function is markedly different than other characterized histones in archaea and eukaryotes.

## DATA AVAILABILITY

Raw sequencing data (ChIP-seq, RNA-seq, and Δ*hpyA* whole genome resequencing) are freely available through the National Center for Biotechnology Information (NCBI) accession PRJNA 703048. ChIP-seq and RNA-seq metadata and additional information can be found on the Gene Expression Omnibus (GEO) through accession GSE182514. All code and input data are available on GitHub at https://github.com/amyschmid/HpyA_codes.

## SUPPLEMENTARY DATA

**Figure S1:** No significant difference in growth rate was observed for WT (blue) and KO (red) strains grown in optimal salt. Y-axis indicates maximum instantaneous growth rate (µ_max_). Crossbars indicate the median of 9 biological replicate trials.

**Figure S2:** OD600 is correlated with CFU/mL across strains and conditions. To calculate colony forming units (CFU), six biological replicate cultures in selected growth phases (exponential and late exponential) were diluted 1 × 10^−4^ to 1 × 10^−6^ depending on the OD600. 100 µL aliquots of each dilution were spread on CM plates. The number of colonies was counted after 7-10 days of growth in a 42°C incubator, and was related to the measured OD600 at the time of plating. In each subpanel, log2 OD600 (x-axis) is plotted against the log10 colony forming units (CFU) / mL (y-axis) for each of the Δ*hpyA* knockout (KO) and Δ*ura3* parent strain (a.k.a. wild type, WT) in optimal salt complete medium (CM) or reduced salt medium (M) as indicated in each plot title. Spearman’s correlation coefficients for OD vs CFU/mL and *p*-values of significance of each correlation are indicated at the top of each plot. Significant correlation between OD and CFU/mL is observed for each strain and condition, indicating strong correspondence between these two measures of growth, and therefore validating spectrophotometric measurements of growth.

**Table S1:** Strains and primers used.

**Table S2:** Raw growth data for parental and Δ*hpyA* cells in optimal and reduced salt (9 biological replicates), measured as optical density (OD600).

**Table S3:** List of peaks obtained by HpyA ChIP-seq; arranged by peak (**S3_simplified**) and by overlap of peak and genomic feature (**S3_full**).

**Table S4:** List of genes differentially expressed in Δ*hpyA* in reduced and optimal salt conditions.

**Table S5**: List of genes in each subcluster obtained by clustering of expression patterns of differentially expressed genes.

**Table S6:** arCOG enrichments for genes nearest the HpyA ChIP-seq peaks, and for genes differentially expressed in Δ*hpyA*

**Table S7**: List of ChIP-seq peaks within 500 bp of differentially expressed genes.

## ACKNOWLEDGEMENTS

We thank current and former Schmid lab members: Cynthia Darnell, Rylee Hackley, Mar Martinez-Pastor, Angie Vreugdenhill, Peter Tonner, Sungmin Hwang, and Preeti Bhanap for technical assistance with experimental methods and analysis, and for comments on the manuscript. Saaz Sakrikar is grateful to his graduate thesis committee members (David Macalpine, Amy Grunden, Richard Brennan, and Raluca Gordan) for mentorship and comments on the manuscript. We thank Deyra Rodriguez and New England Biolabs for providing reagents and advice for rRNA depletion and RNA library preparation. We thank Antoine Hocher and Tobias Warnecke for scientific discussions during the preparation of the manuscript. We thank the Duke Sequencing and Genomic Technologies Core Facility for their technical expertise in generating the sequencing data reported here.

