## Supplementary figures and images for "An archaeal histone-like protein regulates gene expression in response to salt stress"

### Supplementary Figure S1

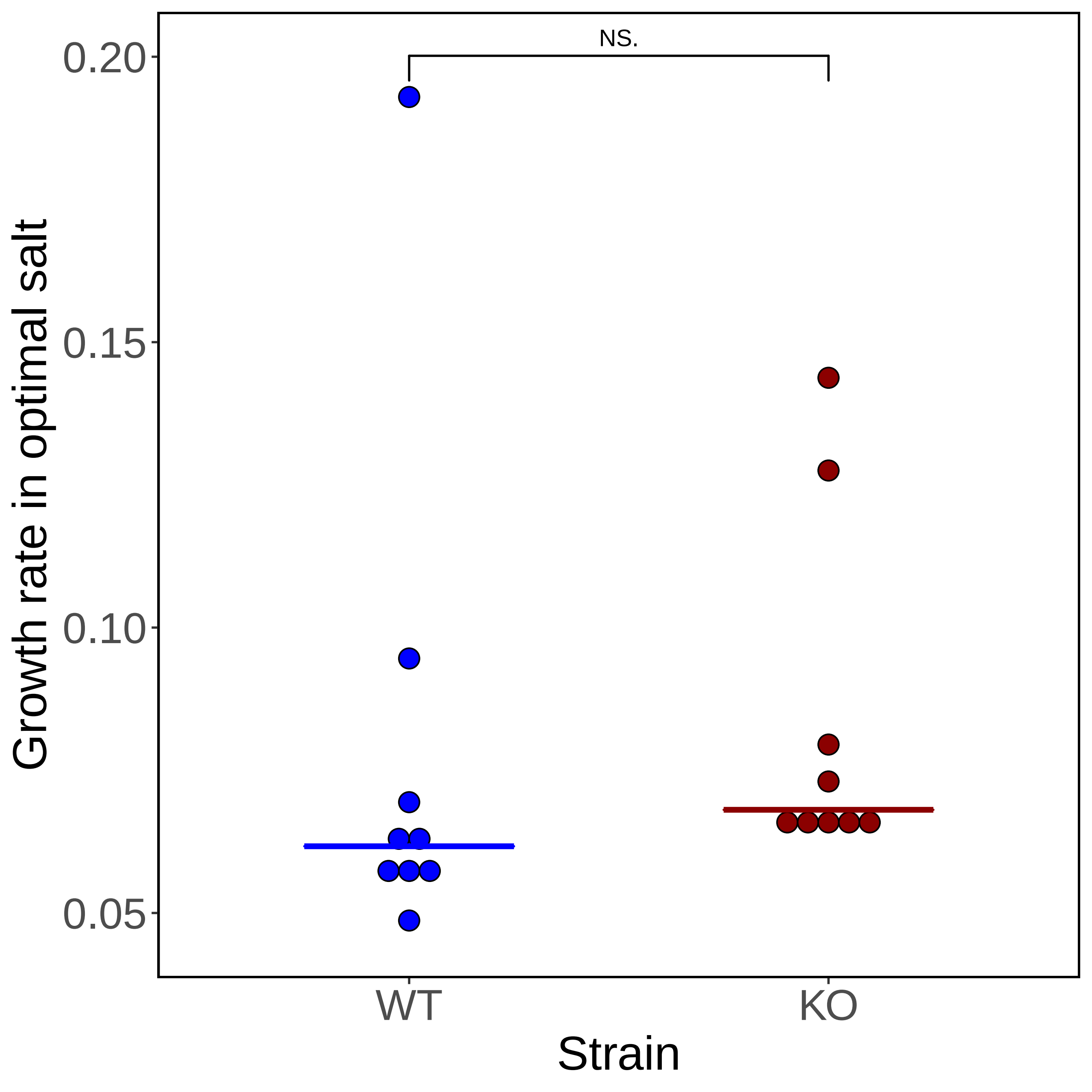

### Supplementary Figure S2

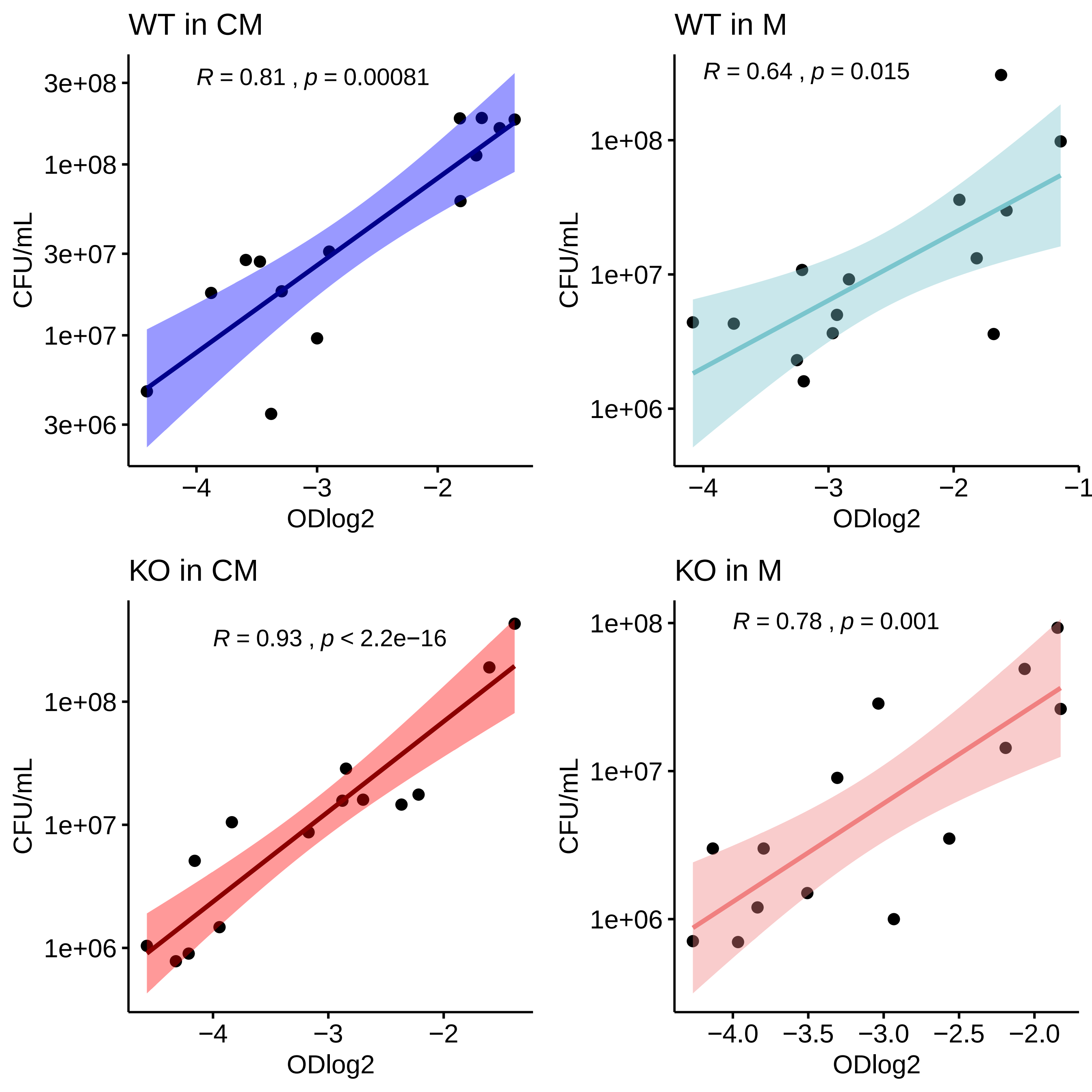
